# Evaluating Human Mutation Databases for ‘Treatability’ Using Personalized Antisense Oligonucleotides

**DOI:** 10.1101/2022.01.04.474998

**Authors:** Swapnil Mittal, Isaac Tang, Joseph G. Gleeson

## Abstract

Genome sequencing in the clinic often allows patients to receive a molecular diagnosis. However, variants are most often evaluated for pathogenicity, neglecting potential “treatability”, and thus often yielding limited clinical benefit. Several collaborative efforts now aim to provide a therapy based upon the genetic variants, even if the drug will benefit only a single patient. Antisense oligonucleotide (ASO) therapies, among others, offer attractive “programmable” and relatively safe platforms for individualized therapy. The landscape of “ASO-treatable” variants is largely uncharted, with new developments emerging for loss-of-function (LOF), haploinsufficient, and gain-of-function (GOF) variants. ASOs can access the genome to target splice-gain variants, poison exons, untranslated/regulatory regions, and naturally-occurring antisense transcripts. Many of these approaches have yet to be proven clinically beneficial, and it is unclear if disease in some patients has progressed past the point where benefit could reasonably be expected. Here we mine public variant databases to identify potential future therapeutic targets. We found that the majority of human pathogenic genetic variants have one or more approaches that could be targeted therapeutically, advantaging the many ways that ASOs can regulate gene expression. The future might see medical teams considering “treatability” when interpreting genome sequencing results, to fully realize benefits for patients.

## Introduction

Next generation sequencing in clinics and in research labs is transforming healthcare, with thousands if not millions of patients per year undergoing exome or genome sequencing to identify genetic causes of disease [1]. Most current emphasis is on the utility of sequencing data to provide an accurate molecular diagnosis relevant to the patient’s clinical features, then using this information to prescribe established therapies to target the disease, sometimes called a ‘top down’ approach. Such an approach has proven beneficial in some patients, but is difficult to scale [2,3], and has largely centered around the type of disease or symptoms. For example, the differential genetic diagnosis for a baby in the NICU with intractable seizures is extensive, but if found to have a biallelic loss of the *ALDH7A1* gene consistent with pyridoxine-dependent epilepsy, then pyridoxine (vitamin B6) can be a lifesaver [2].

An alternative is a ‘bottom-up’ approach, using genetic results to design therapies to treat the variant directly. This alternative has not received much attention because it lacks an established roadmap. Efforts like ClinVar, Online Mendelian Inheritance of Man (OMIM) and the Human Genetic Mutation Database (HGMD) have linked thousands of phenotypes with genetic variations [4–6]. Along with whole genome sequencing (WGS) databases of healthy individuals like gnomAD that track occurrence of genetic variants and along with zygosity, it is now possible to interpret clinical and research WGS with mostly automated pipelines, allowing ever greater confidence in classifying pathogenic variants. With the growth of these databases, and standardization of the interpretation by American College of Medical Genetics (ACMG) and others, the community is gaining confidence in diagnostic clinical sequencing and associated pathogenic mechanisms.

The consideration of *type of variant* rather than the *type of disease* can elucidate a clearer path to treatment. This is especially true for variants previously thought to inactivate gene function, where we now understand a substantial fraction actually activate gene function. Such realization can completely change the therapeutic approach. Classifying amino acid altering variants is particularly challenging in some instances, often requiring experimentation to directly assess the effect of variant on protein function. Clues that some missense variants produce toxic GOF can include clustering in a single domain or even a single amino acid [7]. Thus some dominant alleles, once thought to act through haploinsufficiency, are now instead thought to act through toxic GOF effects [8]. For instance, several tRNA transferase genes with biallelic truncating variants can show infantile encephalopathy where the carriers are unaffected, consistent with LOF and recessive inheritance. But in several of these same genes, a different set of monoallelic variants can show polyneuropathy [9]. While not a critical distinction for clinical geneticists, this can profoundly impact consideration of treatability of the variant, as LOF genes are often approached by restoring healthy copies whereas GOF genes are approached by targeting the diseased allele, gene or protein for degradation (Fig. 1).

**Fig. 1.**
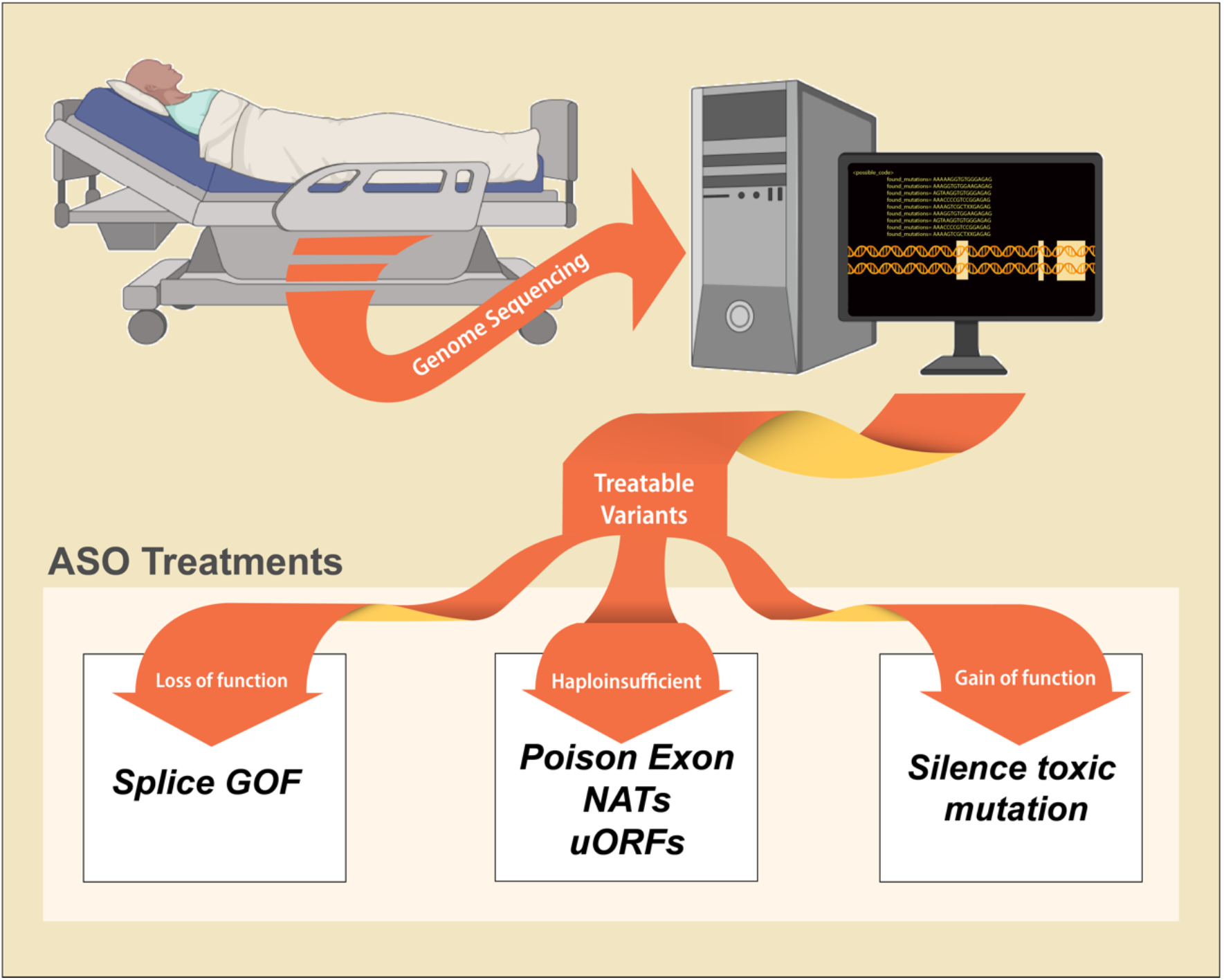
Patient genome sequences analyzed for the potential ‘treatability’ of individual variants. Future methods of interpretation could flag variants for the medical team where there is a special opportunity for potential treatment with approaches such as antisense oligonucleotide (ASO). For instance Loss-of-function (LOF) of genes resulting from gain-of-function (GOF) of splicing variants could be approached with ASOs. Loss of haploinsufficient genes could be approached by targeting poison exons, naturally occurring antisense transcripts (NATs), or upstream open reading frames (uORFs) to increase production of protein from a remaining wildtype allele. Toxic Gain-of-function (GOF) of gene could be approached by silencing toxic mutant alleles.

### Focus on the type of variant

This shift in focus from a particular *type of disease* to instead a particular *type of variant* allows an alternative path to consider treatability in this bottom-up approach (Table 1). The effect of a variant on the molecular biology of a gene is independent from the final effect on the functionality of the gene. Thus a distinction between gain-of-molecular-function (GOMF) vs. loss-of-molecular-function (LOMF) of the *variant* compared with GOF vs. LOF of the *gene* categorization allows a more systematic approach when considering treatment. This is necessary because there are some instances where the effects are antithetical, which can have important implications for treatability.

**Table 1.** Variant classification based upon loss-of-molecular-function (LOMF) vs. gain-of-molecular-function (GOMF) variant and LOF vs. GOF of the gene. Box 1: LOMF variants like gene deletions or frameshifts often result in LOF of the gene. These could be approached by ASO by upregulation of the healthy copy of the gene, if one exists, or with gene replacement therapy. Box 2: LOMF variants like loss of stop codons or splice sites could lead to GOF of the gene. These could be ASO treatable by targeting the toxic allele. Box 3: GOMF variants like gain of a cryptic splice site could lead to LOF of the gene. These could be ASO treatable by targeting the cryptic splice site to correct aberrant splicing. Box 4: GOMF variants like missense amino acid changes in a critical domain could lead to toxic GOF of the gene. These could be ASO-treatable by targeting the toxic allele.

|  | LOF gene | GOF gene |
| --- | --- | --- |
| LOMF Variants | ①<br>Gene deletion<br>Frame shift | ②<br>Stop loss<br>Splice loss |
| GOMF Variants | ③<br>Splice GOF | ④<br>Toxic GOF<br>Missense |

1. LOMF variant leading to LOF of the gene: These include gene deletions or most frameshift variants, where function of one or both copies of the gene is lost and are classically approached by gene replacement therapy. However, if one healthy copy remains active, then an alternative approach is to upregulate expression of the healthy copy of the gene by various mechanisms such as targeting of a poison exon with ASO. This approach was recently used to treat an animal model of Dravet syndrome, due to haploinsufficiency of *SCN1A*. By targeting a poison exon in the healthy copy of *SCN1A*, dosage was corrected and lethality was rescued [10].
2. LOMF variant leading to GOF of the gene: These include loss of stop codons, loss of splice-sites or premature truncation variants that can activate the gene. The effect could be to increase the canonical function of the gene, for instance by increasing activity of a kinase on endogenous substrates, or could be through a toxic GOF, sometimes call ‘neomorphic’, for instance by expanding the range of its kinase substrates. Such variants often occur in single-exon genes or in the 3’ end of a gene, thus escaping nonsense mediated decay (NMD). Although relatively rare, examples include penultimate frameshifts in *DVL1* leading to Robinow syndrome [11]. Because there is gene GOF, these can be approached with ASOs by targeting the toxic allele.
3. GOMF variant leading to LOF of the gene: These include gain of splice site (GOS) variants occurring within an intron, or expansion of a tandem repeat, which could lead to non-functional protein. These can be approached by ASO targeting of the deep intronic GOS spice, for instance in the case of *CEP290* related retinitis pigmentosa [12].
4. GOMF variant leading to GOF of the gene: These include missense variants leading to increased function, toxic GOF of the gene, or gene duplications (i.e. increased gene copies). These can be approached by ASO targeting the toxic allele, for example targeting missense alleles in superoxide dismutase (*SOD*) that lead to protein aggregation in Amyotrophic Lateral Sclerosis or targeting a hyperactive sodium channel in the case of *SCN2A*-related epilepsy [13,14].

Most in the biomedical field appreciate that ASOs can silence gene expression in a mechanism similar to RNAi, however because ASOs targeted the pre-mRNA, many additional effects can be co-opted to treat a range of variant types. ASOs function within the nucleus, concurrent with gene transcription, which not only allows for influence on transcription and transcript stability but also on splicing. Therefore, approaches involving modulation of gene regulation, called TANGO (Targeted Augmentation of Nuclear Gene Output), such as alternative splicing, poison exons (PEs), natural antisense transcripts (NATs) and upstream open reading frames (uORFs) are available to ASOs but not RNAi [15]. The development of ASO treatment requires understanding of the mechanism of disease, the cell type mediating the disease, bioavailability of the ASO to the cells and tissue of interest, and understanding of gene dosage during human development, homeostasis, and aging. For instance, a toxic GOMF variant leading to disease could be targeted by an ASO, but if the non-mutated copy of the gene is required for health throughout life, then ASO target should be focused on just the mutant copy, to produce an allele-selective ASO. On the other hand, if the non-mutated copy is not required for health, then a non-allele-selective ASO may be a more straightforward and ultimately more successful path.

Here we consider several potential mechanisms by which variants could be treated with ASOs and examine several recent breakthroughs that have begun to exploit these approaches. We also examine published databases of variant and their associated genes to infer their treatability by the discussed methods. To be clear, targeting the variant with an ASO, even when leading to the desired effect on target, does not necessarily mean that the patient will show clinical improvement. For instance, the disease may be too far advanced, or the disease progression too slow, to demonstrate clinical benefit. Or certain clinical aspects might improve, while others remain stagnant. By correcting the genetic lesion with ASOs, the field of medicine will learn which diseases and disease classes show clinical improvement.

With this in mind, we profile public variant databases to estimate the fraction of variants that could be approached with ASOs. Such ‘bottom-up’ approaches are now feasible, due in part to the scalable, adaptable nature of ASO design and the well-established safety profiles. So far the experience is very limited [16]. It seems unlikely in the near future that these therapies will be profitable for manufacturers; it has thus, fallen to academic, industry, government, and foundation partnerships to advance the scope of these potentially life-altering therapies.

### Characterizing variants from lens of treatability

We propose categorizing the type of a variant is the first step in considering treatability. This can often be determined from published variant databases such as ClinVar. Of the 1 million entries in ClinVar, approximately 88% are single nucleotide variants (SNVs). Of the approximately 15% categorized as pathogenic or likely pathogenic, the proportion of SNVs drops to just 58%, while deletions (24%) and duplications (9%) make up the majority of the remainder [4] (Fig. S1). Complete gene deletions, frameshift and loss of canonical splice sites are almost always categorized as LOF of the gene, but care is warranted, since some may produce toxic GOF effects if the gene product escapes NMD. For example, truncation of exon 11 of *LAMNA* transcript produces the aberrant ‘progerin’ protein, a cause of Hutchinson–Gilford progeria syndrome, for which ASO targeting shows some benefit [17].

Categorizing a missense variant based upon the likelihood of producing GOF or LOF of the gene can be challenging, and in some instances will require functional validation. For instance, variants appearing in the Catalogue Of Somatic Mutations In Cancer (COSMIC) database are often assumed to produce gene GOF, but COSMIC also contains many LOMF variants in tumor suppressor genes [18]. Emerging efforts like high-throughput functional screens to evaluate every possible non-synonymous missense variants could speed the process [19], but even these may miss toxic GOF if only one criteria of gene function is assessed, for instance effects of variants on protein enzymatic activity. Alternatively, algorithms are emerging to classify individual missense variants as GOF or LOF of gene, for instance the GOF/LOF Database uses a natural language processing of abstracts referenced in ClinVar [20]. The 4785 allele IDs profiled in the GOF/LOF Database represent less than 0.1% of all ClinVar variants, and just 0.3% of all ClinVar pathogenic variants, because the vast majority of ClinVar variants lack sufficient information for classification. Of the variants profiled, 11% (550) were GOF and 89% (4407) were LOF (Fig. S1, Table S1). Thus 77% (3698) of all variants in the GOF/LOF Database are labelled as pathogenic in ClinVar compared with just 15% (157,385 of 1,076,505) of total variants (Fig. S1). However, of ClinVar’s own annotated variants, 680,039 (89%) were predicted to be LOF, which is comparable with the results of the GOF/LOF database. Together these efforts could benefit treatability-predictions.

In the cases of novel variants, where there is little existing information on effect, the type of variant might predict GOF vs. LOF on the gene. For instance, although SNVs make up the majority of reportable variants, there are substantial differences in the relative distribution of mutation type between the LOF and GOF variants. From the GOF/LOF Database, 12% of LOF variants were from deletions, with another 5% from duplications, but these two combined make up less than 4% of the GOF variants, over 95% of which are SNVs. There could result from inadvertent biases in these classifications, tending to assume LOF over GOF. This may lead to corrections in the future as functional data emerges, but clearly not all variant types contribute equally to GOF and LOF effects (Fig. S1).

### LOMF variants causing LOF of gene

Variants causing LOF of haploinsufficient genes (i.e. LOF of one copy of a gene) can be approached either by upregulating the unaffected allele or ortholog if one exists. ASOs can be used to increase expression levels via at least three mechanisms: by promoting poison exon (PE) skipping, by blocking translation of non-canonical open reading frames (uORFs) or translation inhibitory elements, and by blocking regulatory naturally occurring antisense transcripts (NATs). These techniques, grouped together as Targeted Augmentation of Nuclear Gene Output (TANGO), are gaining recognition as viable approaches to therapy [15].

#### Poison Exons (PEs)

PEs are natural, highly conserved, alternatively spliced (i.e. cassette) exons that contain a premature termination codon leading to nonsense mediated decay (NMD) of the transcript if spliced in (Fig. 2A). For example, alternative choice of inclusion or exclusion of a PE in the *FLNA* transcript is a factor in maintaining apical progenitor cell fate in neural progenitors during cerebellar development [21]. Since their initial description, several systematic investigations of alternative splicing have uncovered an abundance of PEs for many genes in the genome, shedding light on their functions. Computational mapping of expressed sequence tags to the human genome reveals that ∼35% of alternative splicing events and 23% of all isoforms contain PEs [22]. Consequently, blocking splicing to a PE can be used to upregulate functional transcript level, whereas prompting splicing can downregulate the level of stable transcript. Proof of concept studies targeting PE splice sites with ASOs for the genes *PCCA* and *SYNGAP1* yielded a 13 to 14-fold reduction of the level of PE-containing transcript and a twofold increase in productive mRNA expression [23]. This data suggests that roughly half of the transcripts for these genes are spliced in a ‘non-productive’ fashion, and that an ASO could potentially double the amount of protein yielded from a single healthy gene copy. Such a rescue might restore protein levels to a degree that could rescue haploinsufficient gene loss. Another example is an ASO targeting PE splicing in *SCN1A*, which was able to increase SCN1A level and largely rescue lethality in a Dravet syndrome mouse model. The treatment resulted in an increased survival in the mouse from 20% to a remarkable 90% [10]. Clinical trials utilizing this approach are now enrolling patients (NCT04442295).

**Fig. 2.**
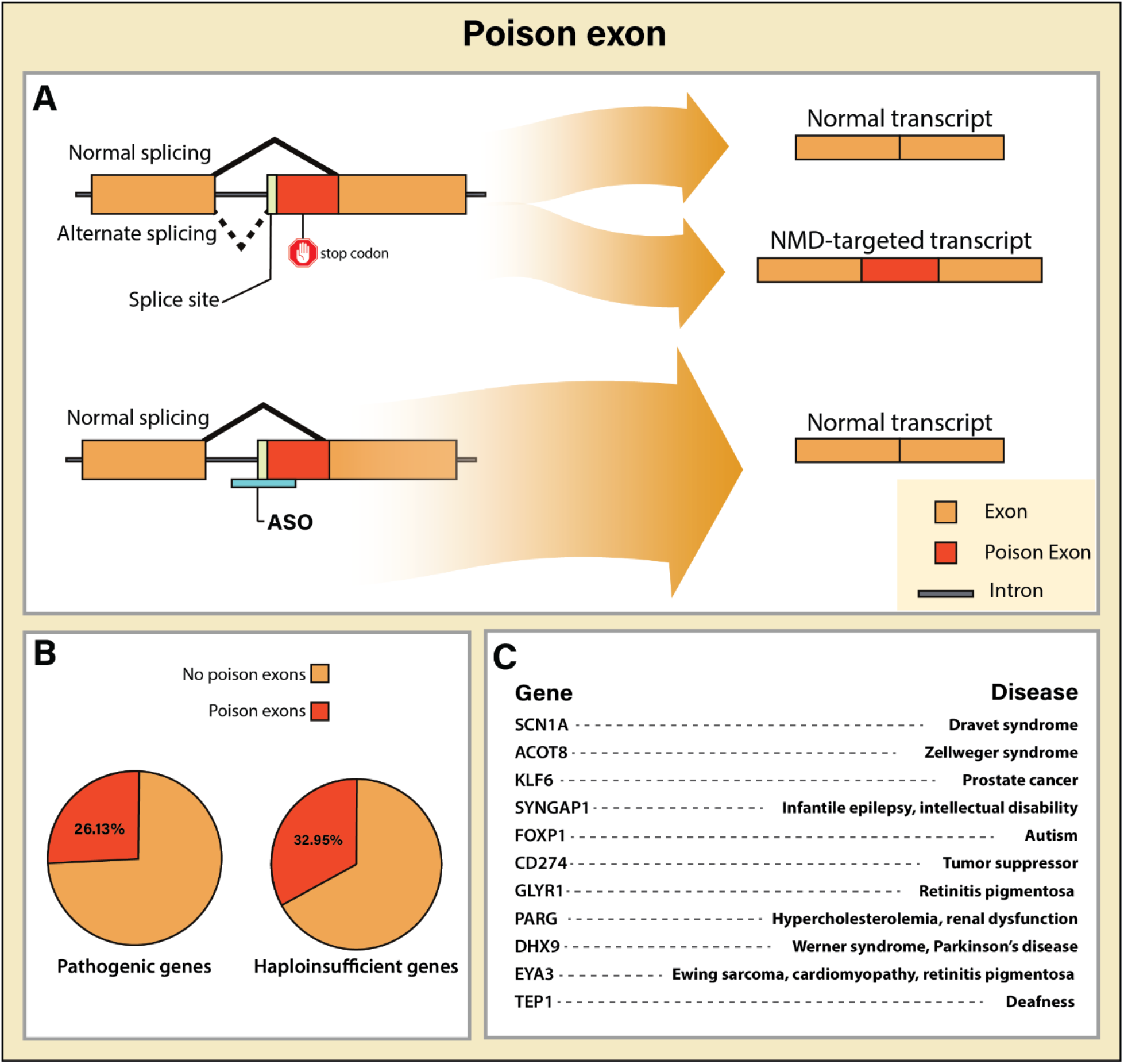
Targeting ‘poison exons’ with ASOs can increase protein production when one copy of a gene is mutated. (A) Poison exons, when incorporated into a transcript, lead to nonsense mediated decay (NMD) targeted transcripts. By ‘cloaking’ the splice acceptor with an ASO (teal), splicing to the downstream productive exon can be restored, and with it, production of protein that may compensate for haploinsufficiency. B) Approximately 26.13% of human genes with pathogenic ClinVar variants and 32.95% of haploinsufficient and likely haploinsufficient genes from ClinGen contain one or more poison exons that could be targeted by ASO. (C) Examples of genes and associated diseases harboring poison exons.

To examine the abundance of PEs in the genome, we collated all spliced exons predicted to result in NMD in the GENCODE and RefSeq databases that did not appear in any protein coding transcript of the same gene (See methods). We found a total of 40,319 potential PEs from a total of 4472 unique genes containing PEs, because genes often have multiple independent PEs, allowing for potentially multiple targets when designing ASOs against PEs (Fig. S2). To determine which specific genes might benefit from a PE ASO approach, we looked at two well established methods for LOF relationships: haploinsufficiency scores in ClinGen and pLI scores in gnomAD (Fig. S1). Of the genes containing PEs, 1589 were classified as pathogenic, and 115 had a haploinsufficiency score of 2 or greater (i.e. haploinsufficient or likely haploinsufficient) in ClinGen [24]. In addition, 831 (i.e. 27%) of genes from gnomAD with a pLI score ≥ 0.9 (i.e. highly intolerant of LOMF variants) showed evidence of at least one PEs (Fig. 2B). This suggests that there are a wealth of disease genes where patients might benefit from ASOs targeting PEs.

Although PEs provide a potential target to increase the expression level of transcripts, separating those in which ASO treatment would substantially increase protein levels could be challenging. Transcripts incorporating PEs can rapidly undergo NMD, making them difficult to detect by conventional RNA sequencing methods, and thus the percent of transcripts in which these PE are ‘spliced in’ (i.e. Percent Spliced In or PSI) may be underestimated. Inhibiting nonsense mediated decay (NMD) can prevent their degradation and produce more accurate PSI assessments. New techniques capable of detecting nascent RNA transcripts can also help overcome this challenge. Global run-on sequencing (GRO-seq), for example, captures RNA during active transcription (i.e. engaged with RNA Pol-II), with the potential to identify PEs [25]. This method might also help identify transcripts targeted for NMD. It is also important to consider that spicing patterns may differ based upon cell type or cell state, so assessing these patterns in functionally relevant cells could be critical.

#### Naturally occurring antisense transcripts (NATs) and other non-coding RNAs

NATs are a subset of non-coding RNAs (ncRNAs) that are transcribed in the opposite direction (antisense or complementary) to one or more sense transcripts. NATs can regulate sense transcripts by RNA-interference (RNAi) through RNaseH mediated decay. Therapeutically, molecules designed to target NATs such as ASOs, or single stranded siRNAs have been termed as ‘antagoNATs’ (Fig. 3A), and can be used to upregulate the expression of cognate sense transcripts, such as for the *BDNF NAT* (i.e. *BDNF-AS*). Downregulation of *BDNF-AS* by siRNA targeting resulted in 2 to 7 fold upregulation of BDNF mRNA both *in vitro* and *in vivo* [26].

**Fig. 3.**
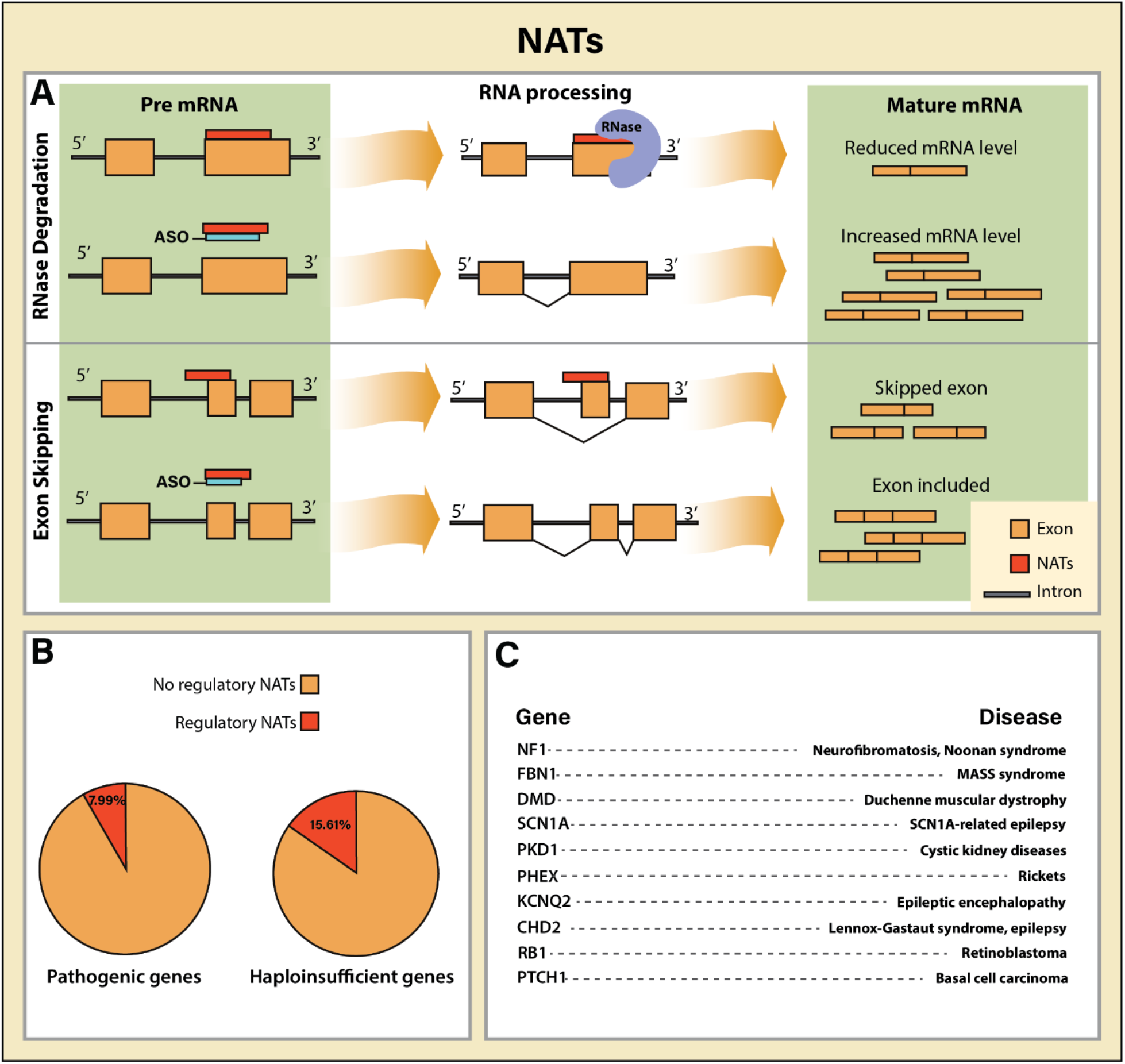
Targeting ‘natural antisense transcripts’ (NATs) may restore protein levels in haploinsufficient conditions. (A) NATs (red) may decrease gene expression, for example by recruiting RNases (purple). ASO targeting NATs can increase transcript levels by inhibiting NAT-mediated repressive effects. NATs may also induce exon skipping or have other repressive effects, which can be disrupted using ASOs to increase certain splice isoforms. (B) Approximately 7.99% of human genes with pathogenic ClinVar variants and 15.61% of haploinsufficient and likely haploinsufficient genes from ClinGen have regulatory NATs that can be assessed for effectiveness of ASO targeting. (C) Examples of genes and associated diseases harboring NATs.

NATs include long noncoding RNAs (lncRNAs), but only recently has the potential for lncRNAs targeting in a therapeutic setting been explored. The *UBE3A-antisense* (i.e. *UBE3A-ATS*) silences the paternally-inherited *UBE3A* allele in a mechanism termed ‘imprinting’. This process contributes to Angelman syndrome in the setting of a maternally inherited LOMF variant. Silencing *UBE3A-ATS* with ASOs can restore *UBE3A* expression and ameliorate some cognitive deficits associated with the disease, in a fashion similar to CRISPR-targeting *UBE3A-ATS* [27,28]. This approach is now in clinical trials.

Another example is the *Chaserr* lncRNA, which represses the chromatin remodeler Chromodomain helicase DNA binding protein 2 (CHD2) in cis. CHD2 haploinsufficiency is associated with a form of developmental epileptic encephalopathy. Knockout of *Chaserr* results in up to a 6-fold increase in CHD2 levels, suggesting that ASO targeting *Chaserr* could be an avenue for treatment in CHD2 disease [29].

NATs can regulate transcription through multiple mechanisms, including direct engagement of a sense transcript through Watson-Crick base pairing, co-transcriptional recruitment of activators or repressors, or long-range genomic-imprinting effects, and for this reason, effect of ASO NAT targeting on sense transcript and protein levels can be difficult to predict, and often requires direct experimentation to measure the direction and effect size on levels of protein [30]. For example, expression of *ZEB2* is regulated by a NAT that targets a splice site in the *ZEB2* 5’-UTR, which also contains an internal ribosomal entry site (IRES). Binding of the NAT prevents splicing of the UTR into the transcript, leading to the retention of the IRES, and subsequent increase in the levels of ZEB2 protein [31].

To evaluate the abundance of NATs in the genome, we compiled a list of all transcripts annotated as ‘antisense’ within the Ensembl database, yielding 12,963 transcripts. These transcripts ranged in length from 54 bp to 92 kbp (i.e. nearly 2000-fold range) [32]. These NATs showed matches to 3844 unique complementary genes. Of these, 495 genes are associated with pathogenic variants in ClinVar (Fig. S2). Most NATs have a single isoform, but some demonstrate alternative splicing. One remarkable outlier is *PCBP1-AS1*, which has 239 annotated splice isoforms, ranging from 208 to 3,908 bp in length. Although *PCBP1-AS1* has been implicated in hepatocellular carcinoma, the effect of this alternative splicing has not been explored [33]. Of all haploinsufficient genes in ClinGen, 54 (16%) have one or more antisense transcripts. Of all genes in gnomAD with a pLI score ≥0.9, 338 (15%) genes had one or more annotated NATs (Fig. 3B). Unfortunately, the manner in which these NATs were annotated can currently only be determine base upon sequence complementarity, and as such, potential regulatory functions cannot be easily predicted. Likewise, ASOs against NATs must be carefully validated because the function of NATs cannot be easily predicted solely from sequence homology, and because NATs, like their sense counterparts, may be subject to alternative splicing. Establishing which diseases might be treated by targeting NATs will likely require cell-based screens to understand these relationships and mechanisms of action.

#### Upstream open reading frames (uORFs) and translation inhibitor elements (TIEs)

uORFs are alternative open reading frames (ORFs) that occur upstream (i.e. 5’) of a canonical main ORF (mORF) and may also encode proteins themselves [34], or may downregulate translation of the mORF by ‘stealing’ ribosomes from the mORF and stalling the scanning ribosome as it reads the upstream stop codon. By this mechanism uORFs can cause up to 80% reduction in mORF protein levels [35] (Fig. 4A). Interestingly, variants with new upstream start codons or disrupting stop sites of existing uORFs are under strong negative selection, emphasizing the importance of their regulatory function [36].

**Fig. 4.**
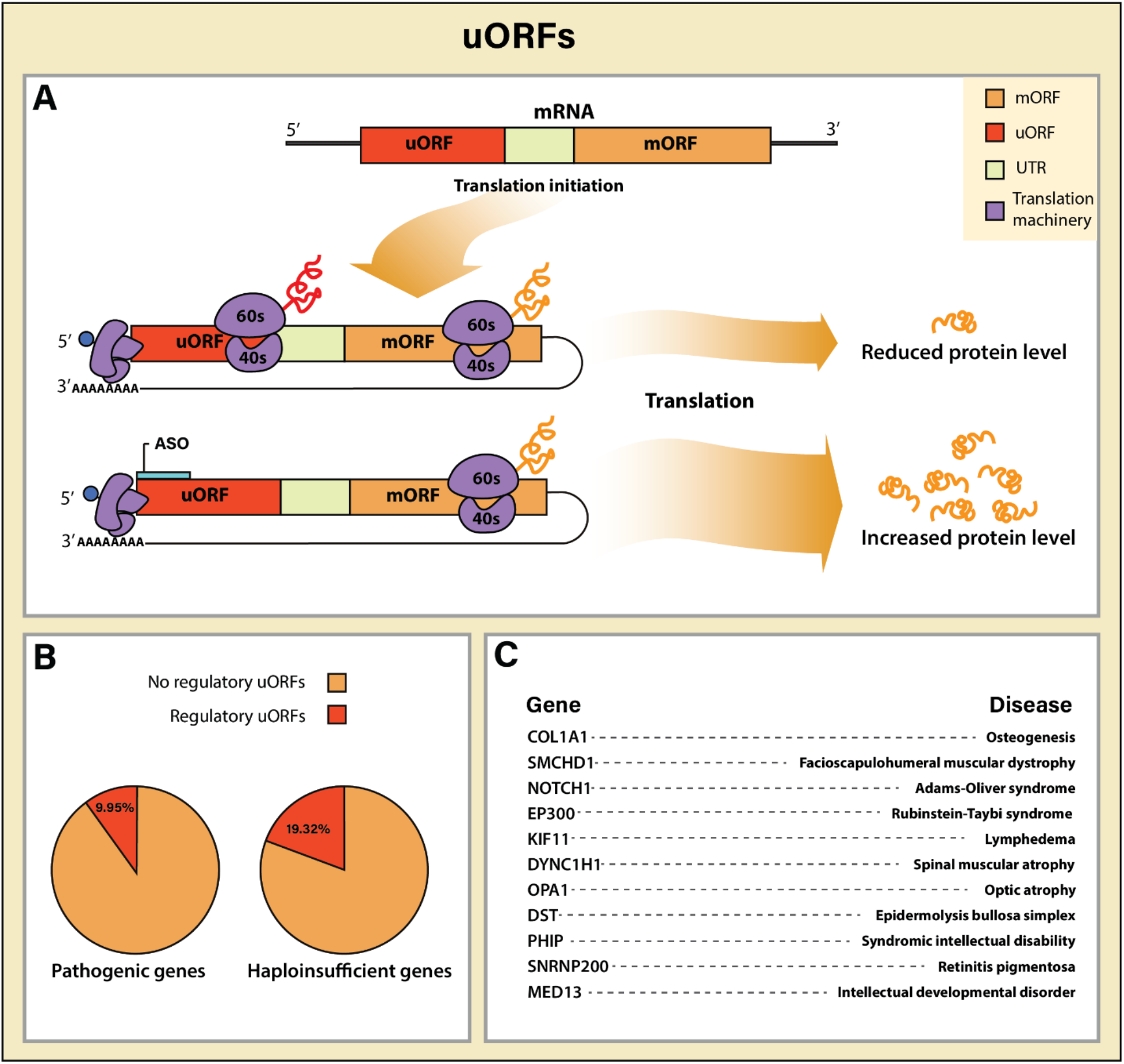
Targeting ‘upstream open reading frames’ (uORFs) by ASOs in haploinsufficient conditions can increase protein levels. (A) Regulatory uORFs reduce protein levels by sequestering the translation machinery (purple) away from main open reading frames (mORFs). Blocking translation initiation of uORF by ASO (teal) allows for higher rates of translation of the mORF compensating for the LOF of the other copy. (B) Approximately 9.95% of human genes with pathogenic ClinVar variants and 19.32% of haploinsufficient and likely haploinsufficient genes from ClinGen contain one or more uORF that could be targeted by ASO to increase protein levels for haploinsufficient diseases. (C) Examples of genes and associated diseases harboring uORF.

ASO targeting of uORFs has been suggested as a mechanism to upregulate mORF protein production. ASOs targeting the *LRPPRC* uORF demonstrated a remarkable 80% increased LRPPRC protein levels without impacting mRNA levels [37]. uORFs are cell-state dependent regulators of gene expression that allow for dynamic changes in protein production during state changes like stress and often act as translational switches [38], mediated in part by cell state induced phosphorylation of eukaryotic initiation factors (eIFs), facilitating bypass of the uORF by the ribosome to promote translation of the mORF [39]. Therefore the effect of ASO targeting of uORFs may be dependent upon cell state [37].

While the majority of putative polypeptides encoded by uORFs are probably non-functional, a fraction appear to contribute to cellular fitness [40], and thus ASO targeting uORFs should be done with caution. Moreover, the affinity of the ASO for the uORF must be high enough to obstruct usage of the uORF, but low enough to allow for its removal by ribosome-associated helicase to allow mORF translation. This property is predicted to result in bell-shaped dose-response curves [37,40].

While uORFs can be predicted from sequence analysis, identifying which uORFs regulate protein levels derived from their cognate mORF requires complementary methods such as Ribo-Seq, in which ribosome complexes are isolated. While this method allows for the identification of novel ORFs, identifying potential fusion polypeptides have proven difficult [41]. We used a database generated from the ORF-RATER algorithm, which relies on linear regression to identify novel and annotated translated coding sequences (CDSs), including uORFs [40]. From this database, 2606 uORFs were identified in 1934 unique genes. Of these genes, 605 (31%) are associated with pathogenic variants in ClinVar, 67 (19%) with haploinsufficient genes in ClinGen, and 523 (17%) with pLI scores ≥0.9 in gnomAD (Fig. 4B). Thus targeting uORFs with ASOs could find a place in therapy for haploinsufficient genes.

TIEs are stem-loop structures present in the 5’ UTR of many mRNAs that either inhibit the assembly of the translation machinery or cause ribosomes to stall. Unlike uORFs, TIEs can be extremely challenging to identify since their effect on translation relies on the secondary structure of the mRNA rather than primary sequence. Regardless, ASOs applied to these sites can also significantly increase ribosome engagement at the mORF site, and thereby boost protein levels [42].

To summarize the potential to treat various LOMF variants leading to LOF of the gene, approximately 55% of haploinsufficient genes from ClinGen and 46% of genes with a pLI score ≥0.9 could potentially be approached with ASO targeting using TANGO (i.e. either NATs, PEs or uORFs) mechanisms (Fig. S2, Table S2). Thus the future is bright for ASO approaches to treat instances of LOMF variant LOF gene.

### GOMF variants causing LOF of gene

#### Silencing toxic gain of splicing variants

A GOMF variant adds functionality to the molecule on which it occurs. The most common and impactful are SNVs that create novel splice acceptor or donor sequences. Sometimes called ‘gain of splice’ (GOS) or cryptic splice sites, these variants ‘trap’ gene splicing, and lead to non-productive mRNAs (Fig. 5A). The general outcome of such an event is to inactivate the gene (i.e. LOF gene) that may cause diseases displaying dominant haploinsufficient, or recessive inheritance. For example, 3% of cystic fibrosis cases 8 are caused by aberrant splicing of CFTR mRNA due to the creation of cryptic splice sites. The mutant splice isoform carries premature stop codons which induce NMD, reducing the expression of the mRNA [43]. Blocking these gain-of-splice-site (GOS) variants using ASOs could be an effective strategy to force expression back to the canonical splice-isoforms. For recessive diseases due to compound heterozygosity, even if one of the variants falls into this class, ASO targeting should be corrective. For instance, the most common genetic form for Leber congenital amaurosis in European descendants is a deep intronic GOS variant in CEP290, seen as homozygous or compound heterozygous [44]. An ASO targeting the intronic variant was able to restore splicing and rescue photoreceptor development in retinal organoids [45].

**Fig. 5.**
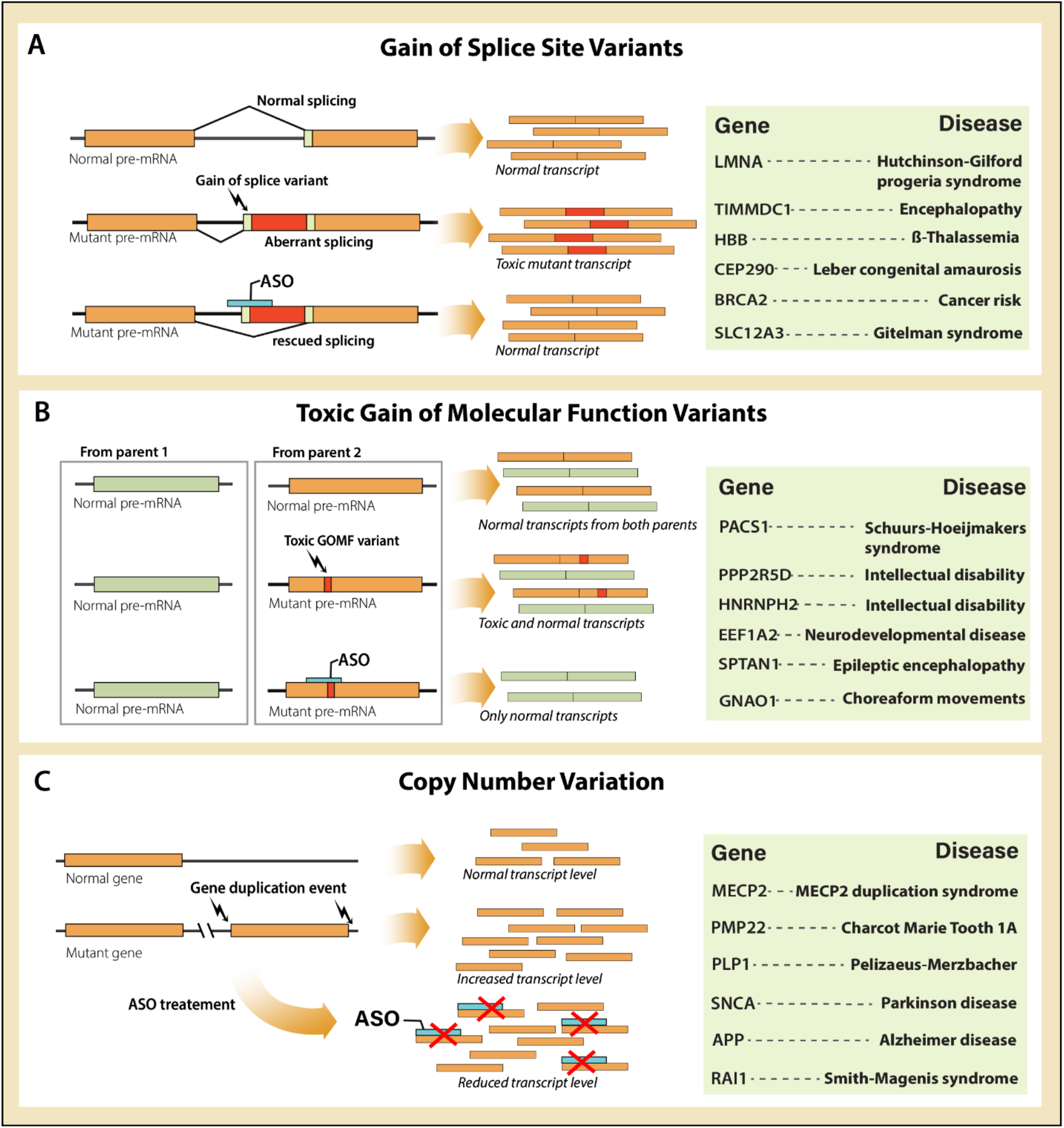
Gain-of-molecular-function (GOMF) variants can be targeted through ASOs and rescue the disease phenotype. (A) Gain of splice sites (GOS) can result in inclusion of intronic regions (red) which can lead to disease. ASOs (Teal) can bind to the mutant splice site and block the toxic splicing. The table shows notable genes with splice gain variants. (B) Toxic GOMF variants can silenced by a ASO specific to the mutant allele (orange) with no significant effect to the normal allele (green). The table shows notable genes with diseases caused by toxic GOMF variants. (C) Copy number variation of genes can cause disease due to increased level of gene products. ASOs can be used to reduce the level of transcript to normal levels in a dose dependent manner. The table lists notable genes with copy number variation related diseases.

Subsequently, 11 patients treated with the intravitreal ASO ‘Sepofarsen’ showed variable improvement in visual function [46]. It remains unknown if greater benefit was not observed because of lack of engagement of the target by the ASO, because of lack of rebound expression of CEP290, or because the disease was too far progressed to demonstrate benefit.

A critical need is the development of comprehensive databases of disease-related GOS variants. Attempts such as DBASS3 and DBASS5 [47] record all instances of aberrant mRNA due to de novo variants, however predicting the consequence of noncoding variants often requires analysis of RNA. Machine learning algorithms such as SpliceAI could help aid in the search of disease causing GOS variants. There is also a lack of computational methods to predict specificity and efficiency of a particular ASO in regulating the target locus. This is partly due to complex interactions between ASO and mRNA that determine binding efficiency and effect on protein production. High throughput ASO screening can partially circumvent this problem by selecting the most effective candidate ASOs from large pools. For instance, a library of 198 ASOs representing different sequences and backbone chemistry was generated to target the aberrant splicing of LMNA pre-mRNA. A gain of 5’ splice acceptor variant in this gene leads to formation of progerin mRNA, a cause of Hutchinson–Gilford progeria syndrome. ASOs were then screened against the pre-mRNA to identify those that could most effectively force splicing of healthy lamin A and lamin C isoforms. The top-performing ASO reduced progerin transcript levels by 90% and increased mean survival of progerin mice by 100 days [17].

### GOMF variants causing GOF of gene

#### Silencing toxic gain of molecular function (GOMF) variants

One would think that GOMF variants leading to a GOF or toxic GOF of the gene would be rare, but accumulation of data in the field of human genetics has highlighted this as an important and often overlooked contribution to pathogenesis. This is especially true for *de novo* missense (DNM) variants. Previously it was thought that most DNMs led to partial or complete gene LOF, but recent advances have shown that toxic GOFs are more common than expected. Many such DNM-GOMF variants are highly spatially clustered within the protein, often occurring in autoregulatory domains, and can have dramatic impact on protein function [7]. These protein domains often show constrained variability, which suggests a strong purifying selection that foreshadow severe developmental phenotypes or embryonic lethality when mutated [49].

Ideally a non-allele specific ASO could be used to target a GOMF variants if there were no adverse consequences of knockdown of transcript from both copies of the gene (i.e. the mutant and the wildtype copy). But in practice it is often difficult to determine whether knockdown of the healthy copy is functionally important. Even if homozygous mouse knockouts shows no phenotype, and where LOMF variants are known to exist in humans, it would be risky to completely or almost completely inactivate a human gene, even in the setting of one copy containing a toxic GOMF variant. A confounding finding is that some genes show overlapping phenotypes produced from a LOMF and GOMF variant. For instance, certain DNM-GOMF variants in the BAF complex component *ACTL6B* lead to encephalopathy. Yet in this same gene, presence of biallelic LOMF variants also lead to encephalopathy [8]. For this reason, a non-allele-selective approach would be predicted to inactivate both copies, and produce no clinical benefit. But if sufficient allele-selectivity could be achieved with an ASO, it should be possible to reduce levels of the toxic allele while leaving the wildtype allele intact, to yield potential clinical benefit. Similarly, DNM-GOMF alleles in Golgi *PACS1* lead to encephalopathy [50], where mouse biallelic knockouts are healthy. Would non-allele selective targeting of mutant *PACS1* be prudent?

The challenge lies in creating allele-selective ASOs which knockdown of a GOMF variant since with minimum off target effects (Fig. 5B). The process of identifying allele-selective ASOs is greatly aided by identifying additional SNPs phased to the toxic GOMF haplotype to which ASOs can be screened. Long-read sequencing could help phase distal SNPs or INDELs, while avoiding variants that are present on both haplotypes. Because a polymorphism occurs approximately every 1000 bp, the average gene size of 50kB should yield approximately 50 SNPs to which allele-selective ASOs could be designed. On the other hand, short genes could prove difficult to design allele-selective ASOs. Future developments may utilize adaptive configuration of the ASO backbone to improve allele selectivity [51].

#### Dose dependent regulation of gene duplications

Gene duplication events can lead to diseases due to dysregulation of gene expression. For instance, *MECP2* duplication syndrome (MDS), seen almost exclusively in males, shows features similar to its female-predominant counterpart, Rett syndrome. ASOs targeting a *MECP2* duplication in a mouse model of MDS not only restore physiological expression levels in an ASO-dose dependent manner but also abolish seizures [52]. Similarly, *PMP22* gene duplications result in Charcot-Marie-Tooth disease type 1A (CMT1A), and even conditions like chromosomal duplication may one day be approachable with ASOs [53](Fig. 5C).

## Conclusions

Bottom-up approaches to treat genetic disease are gaining momentum, with many rare disease family support networks advocating for this therapy. Perhaps this could have been anticipated a decade ago when diagnostic sequencing in the clinic became a reality, but the recent pace has nevertheless been remarkable. ASO treatment for spinal muscular atrophy was the opening bell, and other conditions like Angelman syndrome will be next. Therapies for these two diseases quickly garnered investments from pharmaceutical companies since the same ASOs can be used to treat most patients. Identifying which conditions and haplotypes can be treated with a general purpose ASO, versus which will require a patient-specific solution will be an important undertaking.

Where should physicians, advocacy groups, or patients turn if their disease has no treatment? With the consideration of treatability of the variant, there are now many options for a bottom-up approach. We found that a surprising 55% of haploinsufficient genes could be approached therapeutically, even with our limited understanding of disease pathogenesis. Groups like the n-Lorem Foundation offer the opportunity for nomination of patients with rare genetic disease for treatment, and currently have accepted over 60 patients for personalized ASO production [54]. The n-Lorem Access to Treat Committee conducts genetic and regulatory experiments and meets monthly to consider patient nominations into the protocol. Similar efforts are underway at the n-of-1 Collaboratory in partnership with the Oligonucleotide Therapeutic Society. In parallel, the FDA has issued special guidance for n-of-1 trials, and in response the research community is developing strategies for n-of-1 case designs [55].

In its current form, gene delivery approaches using viral delivery have challenges and opportunities that are quite distinct from ASOs. Access of payload to the desired cells is a challenge, although often the same viral drug can be used irrespective of the underlying variant. The Institute for Life Changing Medicine seeks to produce viral-mediated gene therapy for rare disease. NIH Foundation sponsored Bespoke Gene Therapy Consortium (BGTC) will take advantage of the NIH GMP facility for the same purpose. Many other such approaches, including CRISPR gene targeting and base editing, are also on the horizon.

This grand experiment will test the collaborative nature of researchers, clinicians, patients, philanthropists, and regulatory agencies. At stake could be considered the greatest research medicine has ever undertaken, as we will soon learn which diseases and which genes are amenable to treatment if the genetic variant can be corrected.

## Methods

### General Statistics and Data Analysis

Data analysis was performed using custom scripts in Python 3.8.12 using the Pandas 1.3.3 and Numpy 1.20.3 libraries, available upon request. Plots were generated with the Matplotlib 3.4.3 library and Adobe Illustrator 2021.

### Interrogation of ClinVar, ClinGen, and gnomAD

The ClinVar variant summary freeze 9/27/2021 was downloaded from the ClinVar FTP site, using only variants from assembly GRCh38. We utilized the ClinVar clinical significance terms, grouping ‘pathogenic’ and ‘likely pathogenic’ into ‘pathogenic’, whereas ‘benign’ and ‘likely benign’ were grouped into ‘benign’, and ‘uncertain’ and ‘conflicting interpretation’ were grouped as ‘uncertain’. Variants without ‘clinical significance’ entry were excluded. ClinVar variants containing ‘variation’ or ‘complex’ in the field ‘Types of mutation’ were omitted. When identifying unique genes that contain pathogenic, benign, or unknown variants, multiple genes associated with a single variant were split and separately appended to the gene list. This analysis yielded a total of 1,076,505 variants. The genes that contain pathogenic variants were referred to as ‘pathogenic’ genes.

To compare the following databases with a list of haploinsufficient genes, we downloaded the construction GRCh38 ClinGen haploinsufficiency gene database. Only genes with haploinsufficiency entries were examined. To generate the gene list, genes with haploinsufficiency score of 3 (‘sufficient evidence for haploinsufficiency’) and 2 (‘emerging evidence for haploinsufficiency’) were included, yielding 346 haploinsufficient genes.

To generate a list of genes that are highly intolerant to loss-of-function, we downloaded the gnomAD LOF metrics by gene version 2.1.1., excluding all but pLI scores of greater than or equal to 0.9.

### Identification of toxic GOMF and LOMF variants in ClinVar

The gain of function/loss of function (GOF/LOF) database was downloaded from https://itanlab.shinyapps.io/goflof/, excluding HGMD annotation, from the June 2021 release. GOF/LOF variant entries were mapped to ClinVar variant entries for 4785 allele IDs. Unique gene names from the GOF/LOF database were separately tabulated.

### Identification of Poison Exons in ClinVar (PEs)

Transcripts labelled as NMD in GENCODEv34 and RefSeq 20191206 databases, which did not appear in the protein coding transcripts in the same gene were designated as PEs. Gene name, chromosomal location, and starting and ending coordinates for each of the 40,319 PEs were assembled into a BED file. Genes containing at least one PE were compared with the ClinVar and ClinGen gene lists to determine the pathogenic and haploinsufficient genes that contain PEs.

### Identification of Naturally Occurring Antisense in ClinVar (NATs)

Antisense sequences were acquired through Ensembl and annotated through the R library biomaRt ver 2.5, yielding 12,724 unique transcripts associated with 3852 unique genes annotated as ‘lncRNA’. The unique genes were compiled into a separate list for further comparison with the ClinVar and ClinGen databases to determine the pathogenic and haploinsufficient genes that had NATs. NAT PCBP-AS1 was excluded as an outlier because it contained more than 200 NATs.

### Identification of upstream Open Reading Frames in ClinVar (uORFs)

The database for uORFs was assembled from Riboseq from human iPSCs and human fetal fibroblasts (HFFs) by the Weissman lab [56]. Only protein-coding sequences (CDSs) labelled as ‘upstream’ were designated as uORFs. We then calculated the number of uORFs per gene, and compared to pathogenic and haploinsufficient genes from ClinVar and ClinGen, respectively.

## Online resources

OMIM: https://www.omim.org

gnomAD: https://gnomad.broadinstitute.org

ClinVar: https://www.ncbi.nlm.nih.gov/clinvar/

HGMD: http://www.hgmd.cf.ac.uk

GOF/LOF: https://itanlab.shinyapps.io/goflof/

ANTIcode: http://bioinfo.org.cn/anticode/

SpliceAI: https://spliceailookup.broadinstitute.org/

COSMIC: https://cancer.sanger.ac.uk/cosmic

Weissman uORF: https://pubmed.ncbi.nlm.nih.gov/26638175/

DBASS3 and DBASS5: https://www.dbass.org.uk

## Acknowledgments

The authors wish to thank Jonathan Sebat, Jennifer Friedman, David Dimmock, Charlotte Hobbs, Matthew Bainbridge and members of the Gleeson lab for comments. This work was supported by the Rady Children’s Institute for Genomic Medicine.

## Conflicts of Interest

Dr. Gleeson serves on the Access to Treat Committee and Chief Medical Officer for the non-profit N-Lorem Foundation. Dr. Gleeson consults for Ionis Pharmaceutical, Inc. and Shire Pharmaceutical, Inc.

---

**Figure S1.**
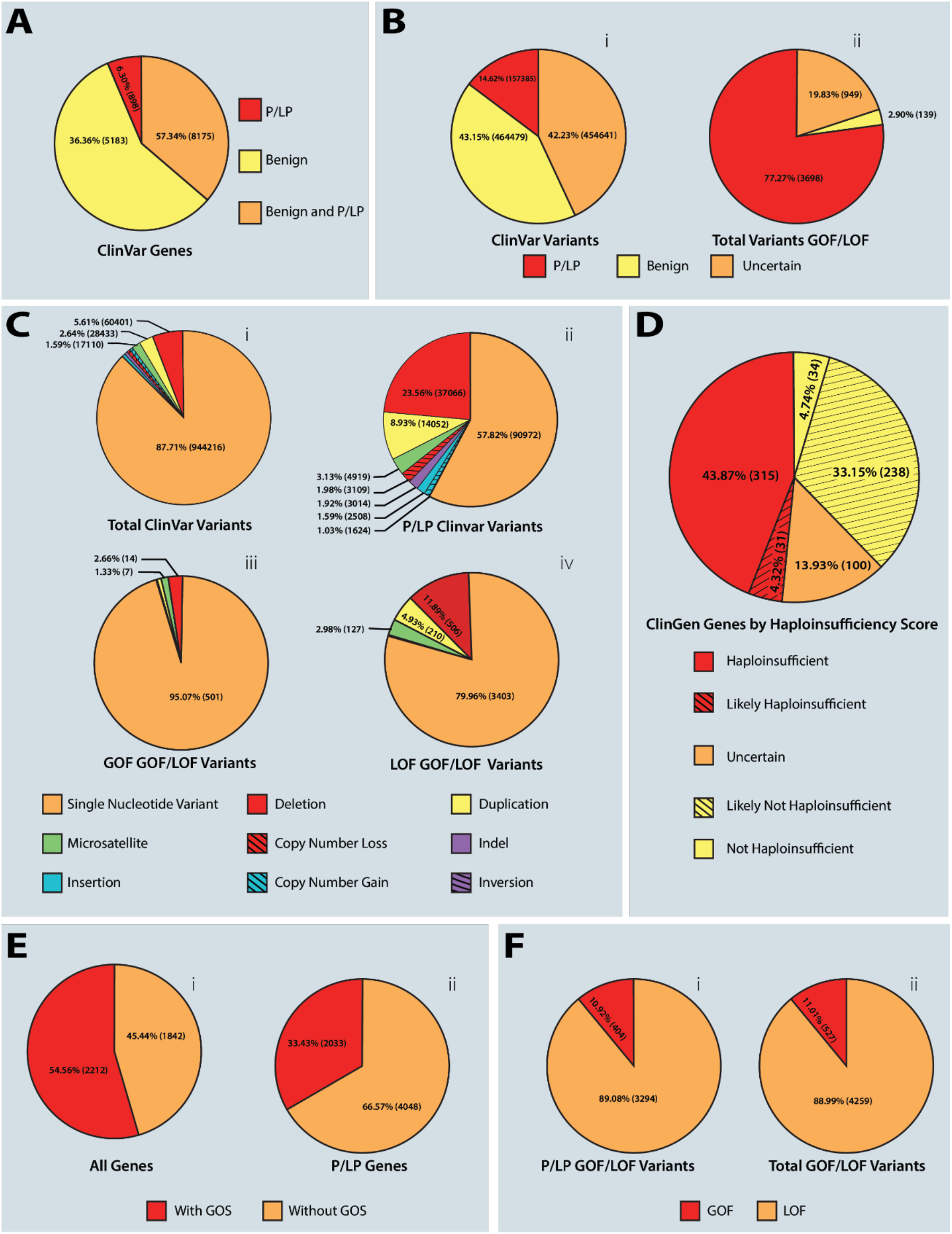
A) Clinical significance of the genes associated with examined ClinVar variants (P/LP: pathogenic or likely pathogenic). B) Clinical significance of variants in ClinVar and GOF/LOF databases. C) Distribution of types of variant in all variants found in ClinVar(i) and GOF/LOF(iii). Distribution of types of variant in all pathogenic variants found in ClinVar (ii) and GOF/LOF (iv). D) Annotated ClinGen genes by haploinsufficiency score. E) ClinVar genes containing gain of splice (GOS) variants (i), and P/LP genes containing GOS variants according to ClinVar (ii). F) LOF vs GOF variants for P/LP(i) and total (ii) variants in GOF/LOF database

**Figure S2.**
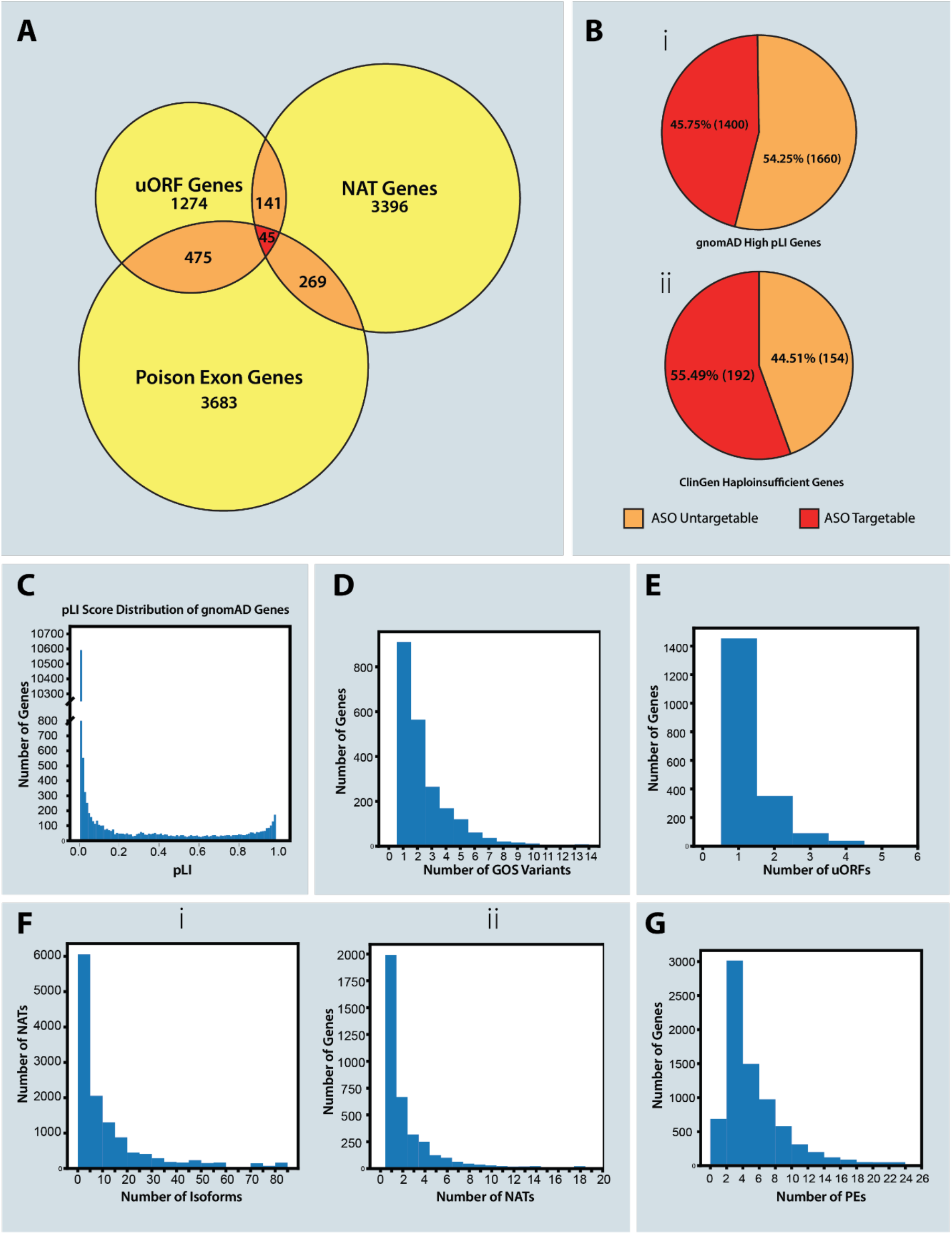
A) Venn diagram of genes containing poison exons (PEs), upstream open reading frames (uORFs), or natural antisense transcripts (NAT). B) Distribution of ‘targetable’ (contains at least one poison exon, uORF, or NAT) genes in the gnomAD high pLI gene list (i), and haploinsufficient genes in ClinGen (ii). C) Distribution of pLI scores of gnomAD genes. D) Number of gain of splice (GOS) variants per gene in GOS database. E) Number of uORFs per gene. F) Number of different splice isoforms per NAT (i) and number of NATs per gene (ii). G) Number of poison exons per gene.

**Suppl Table 1 (First 37 of 4785 entries).**
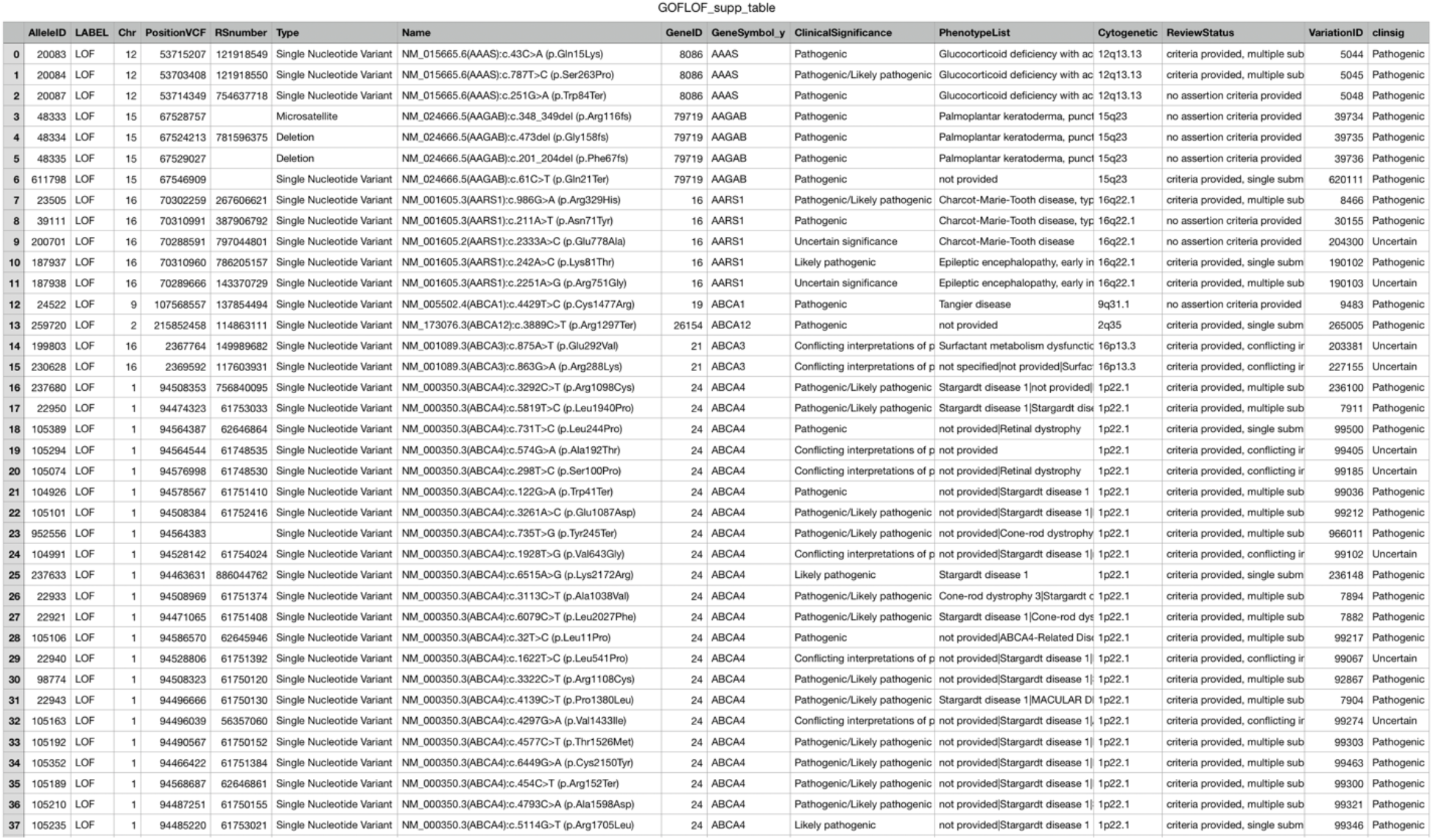
Table of all 4785 allele IDs listed in the GOF/LOF Database, listing their label as either LOF or GOF (column 3), along with the cognate gene, clinical significance and associated phenotype.

**Suppl Table 2 (first 37 of 9282 entries).** Table of all 9282 genes listed in ClinVar with assignment of Pathogenic or Likely Pathogenic variants, compared with the presence of one or more upstream open reading frames (uORFs), naturally occurring antisense transcripts (NATs), or poison exon (PE). Also annotated is whether the gene is listed as ‘haploinsufficient’, whether the gene is ‘pathogenic’ or ‘likely pathogenic’ from ClinVar annotation, whether the gene is known to undergo alternative splicing and the protein Loss of Function Intolerant (pLI) score.

|  | Gene | uORF | NAT | PE | Haploinsufficient (ClinGen) | Pathogenic (ClinVar) | Contains Known Splice Variants | pLI |
| --- | --- | --- | --- | --- | --- | --- | --- | --- |
| 0 | A1BG | N | Y | N | N | N | N | 4.9917e-09 |
| 1 | A2M | Y | Y | N | N | N | Y | 4.5229e-11 |
| 2 | A2ML1 | N | Y | N | N | Y | Y | 1.6109e-34 |
| 3 | A4GALT | Y | N | N | N | Y | N | 7.6668e-06 |
| 4 | AADACL2 | N | Y | N | N | N | N | 1.0604e-08 |
| 5 | AAK1 | Y | N | N | N | N | N | 1.0 |
| 6 | AANAT | N | N | Y | N | N | N | 0.002969 |
| 7 | AARSD1 | N | N | Y | N | N | N | 4.0653e-09 |
| 8 | AASS | N | N | Y | N | Y | N | 2.3101e-12 |
| 9 | AATF | N | N | Y | N | N | N | 2.5958e-12 |
| 10 | AATK | N | N | Y | N | N | N | 0.20634 |
| 11 | ABALON | N | Y | N | N | N | N | N |
| 12 | ABAT | N | N | Y | N | Y | N | 0.0063216 |
| 13 | ABCA10 | N | N | Y | N | N | N | 7.6593e-34 |
| 14 | ABCA13 | N | N | Y | N | Y | N | 8.7753e-114 |
| 15 | ABCA2 | N | N | Y | N | Y | Y | 1.0 |
| 16 | ABCA5 | N | N | Y | N | Y | Y | 4.7803e-27 |
| 17 | ABCA7 | N | N | Y | N | N | Y | 2.1111e-55 |
| 18 | ABCA8 | Y | N | N | N | N | N | 1.6089999999999998e-52 |
| 19 | ABCA9 | N | Y | N | N | N | N | 9.4911e-48 |
| 20 | ABCB11 | N | N | Y | N | Y | Y | 1.5612e-12 |
| 21 | ABCB4 | N | N | Y | N | Y | Y | 2.2138e-07 |
| 22 | ABCB5 | N | N | Y | N | N | N | 1.5071999999999999e-37 |
| 23 | ABCB6 | N | N | Y | N | Y | Y | 8.599900000000001e-19 |
| 24 | ABCB7 | N | N | Y | N | Y | N | 0.99994 |
| 25 | ABCB8 | N | N | Y | N | N | N | 1.9164e-15 |
| 26 | ABCB9 | Y | N | Y | N | N | N | 0.00025164 |
| 27 | ABCC1 | N | N | Y | N | Y | Y | 0.0011727 |
| 28 | ABCC12 | N | N | Y | N | N | N | 4.5659e-49 |
| 29 | ABCC2 | N | N | Y | N | Y | Y | 5.2025e-47 |
| 30 | ABCC3 | N | N | Y | N | N | N | 1.2092e-26 |
| 31 | ABCC4 | N | N | Y | N | N | N | 1.0674e-07 |
| 32 | ABCC5 | Y | Y | Y | N | N | N | 0.0041518 |
| 33 | ABCC6 | N | N | Y | N | Y | Y | 5.6167e-35 |
| 34 | ABCC8 | N | N | Y | N | Y | N | 7.9571e-24 |
| 35 | ABCD4 | N | N | Y | N | Y | N | 1.3364e-12 |
| 36 | ABCE1 | N | N | Y | N | N | N | 0.9738 |
| 37 | ABCF3 | N | N | Y | N | N | N | 1.5114e-05 |

